# Strong correlations between mutation rates and selective constraint in the human genome?

**DOI:** 10.64898/2026.10.08.757506

**Authors:** Gustavo V. Barroso, Aaron P. Ragsdale

## Abstract

The 21st century has seen repeated challenges to the premise that *de novo* mutations occur independently from their fitness effects. So far, these attempts have been dismissed as technical artifacts, restoring confidence in the null hypothesis. Yet the development of high-resolution genomic resources enables investigating the relationship between mutation rate variation and selective constraint with unprecedented detail. Here, we use state-of-the art genomic maps to test whether predicted mutation rates differ within and outside putatively constrained sites in humans. Contrary to the hypothesis that is typically put forward, but in agreement with another recent study, we find that protein-coding genes exhibit higher rates of mutation than the neutral genomic background. Conversely, mutation rates are lower in non-coding elements identified through sequence conservation. This can be attributed either to biases in the estimation of mutation rates in sites under selection, biases in scoring constraint in sites with low rates of mutation, or a genuine signal that would merit further investigation. Regardless of the underlying cause, our findings have broad implications for evolutionary genetics. Several studies rely on accurate mutation maps and identification of selection targets across the genome, which inherently confound each other. Caution is therefore warranted when interpreting results from currently available non-coding scores.

## Introduction

A central tenet of evolutionary biology is that *de novo* mutations occur independently from their fitness effects (Kimura, 1968; Bataillon and Bailey, 2014; Stoltzfus, 2021). Recent studies across diverse model organisms have challenged this premise, advancing the hypothesis that selection has fine-tuned local mutation rates to reduce genetic load. Martincorena et al. (2012) proposed that *Escherichia coli* genes under stronger purifying selection mutate less often, but this pattern was later shown to arise from population-genetic confounding (Chen and Zhang, 2013). In humans, Xia et al. (2020) suggested that “scanning” in testes optimizes germline mutation rates via transcription-coupled repair, but subsequent reanalysis demonstrated that this was an artifact (Liu and Zhang, 2020). More recently, Monroe et al. (2022) reported reduced mutation rates in protein-coding genes of *Arabidopsis thaliana*, but faced criticism on both data-quality and population-genetic grounds (Charlesworth and Jensen, 2023; Liu and Zhang, 2022; Wang et al., 2023; Monroe et al., 2023). Thus, despite repeated attacks, the baseline model of independence between *de novo* mutation rates and selective constraint currently holds. This expectation is rooted in population genetics theory: while there is evidence that second-order selection can plausibly reduce genome-wide mutation rates (Lynch, 2011; Lynch et al., 2016), selection on region-specific mutation is likely too weak to be effective, especially in species with historically low population sizes (Chen and Zhang, 2013).

Yet human somatic cells show evidence of reduced mutation rates in exons due to more efficient recruitment of the mismatch repair machinery (Supek and Lehner, 2015; Frigola et al., 2017). While Rodriguez-Galindo et al. (2020) could not replicate this result in the germline, their analyses involved relatively few mutations, and subtle but biologically meaningful effects may have been missed. Conversely, whether mutation rates differ within and outside non-coding elements such as regulatory or phylogenetically constrained regions, has not been systematically tested. The availability of high-resolution mutation maps in humans (Carlson et al., 2018; Karczewski et al., 2020; Seplyarskiy et al., 2023), each focusing on different aspects of mutation rate variation, together with growing efforts to annotate the human genome, enables revisiting this topic.

We have recently found a signal of covariance between mutation rates and the density of constrained sites in the human genome (Barroso et al., 2026). The covariance was negative for non-coding phastCons (Siepel et al., 2005; Di et al., 2025) and positive for protein-coding genes (which tend to be under stronger selection than non-coding regions in humans). Notably, elevated mutation rates in genes are unexpected if selection favors mutation-rate modifiers that reduce genetic load. Here, we further explore this signal, revealing that predicted mutation rates differ markedly within putatively constrained elements. In particular, the reduction in non-coding elements can reach *>* 30%, even with respect to nearby sites. There are three possible explanations for this pattern: 1) biases in the estimation of mutation rates in sites under selection; 2) biases in scoring constraint in sites with low rates of mutation; 3) genuine differences in mutation rates between neutral and constrained sites.

## Results

### Contrasting mutation rates using publicly available genomic maps

We investigated predicted mutation rates within and outside different annotations in the human genome: protein-coding exons (Harrison et al., 2024), promoters and enhancers (Vu and Ernst, 2022), non-coding phastCons (Siepel et al., 2005), and machine learning scores from `GPN-MSA` (Benegas et al., 2025a). Groups were further divided into constraint classes according to the predicted strength of negative selection (lower corresponding to more deleterious, Methods). We compared mutation rates in constrained elements with the putatively neutral background using the ratio of the average rate within each class, 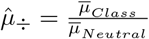.

Our results take advantage of methodological differences among mutation maps. Both Roulette (Seplyarskiy et al., 2023) and Carlson et al. (2018) use polymorphism data from extremely rare variants (ERVs) and fit sophisticated models to separate true mutational processes from the effects of selection. These maps represent the state-of-the-art of how features such as recombination, sequence context (*k*-mers) and replication timing affect human mutation rates (Seplyarskiy et al., 2021). On the other hand, gnomAD rates are based only on trinucleotide context and methylation patterns, serving as a baseline model of mutation rate variation (Karczewski et al., 2020). We show that the three maps often indicate different effect sizes but agree on the shape of the relationships between 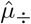 and putative selective constraint.

### Protein-coding sites have higher predicted mutation rates than the neutral background

We found that mutation rates are elevated(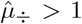, Figure 1A)in protein-coding exons (all maps) and promoters (except Roulette), and are similar to the rest of the genome in enhancers (except gnomAD). At face value, this is surprising since protein-coding exons are among the most constrained sites in the human genome. Hence, under the hypothesis of regional mutation-rate optimization, we would expect them to show lower mutation rates. One possible explanation is that first-order selection for protein function preserves highly-mutable sequence contexts (Chakraborty et al., 1997; Aggarwala and Voight, 2016) shifting *k*-mer distributions away from their neutral steady-state (Oman et al., 2022). Our results suggest that the ensuing increase in exonic mutation rates is substantial, between 10% and 30% depending on the map, consequently increasing mutational load (Ying and Huttley, 2011; Agarwal and Przeworski, 2021). This predicted reduction in population fitness occurs despite the robustness of the genetic code against damaging amino-acid mutations (Haig and Hurst, 1991; Freeland and Hurst, 1998; Loay et al., 2026).

**Figure 1:**
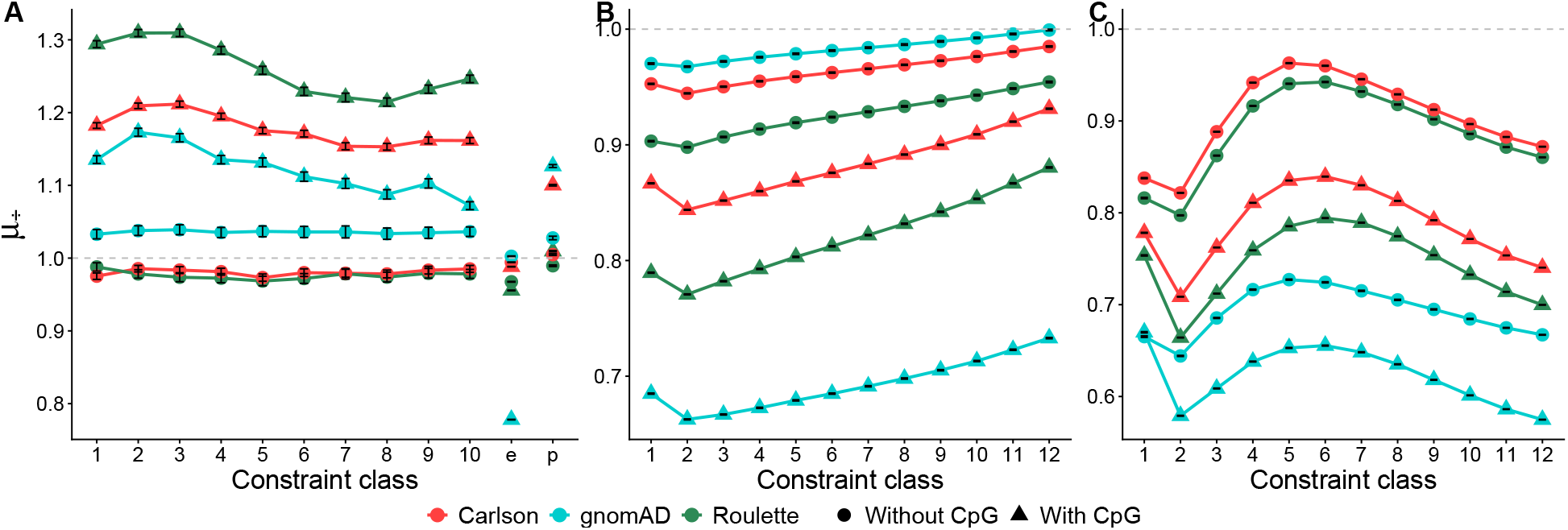
Predicted mutation rates differ within and outside putatively constrained sites across the genome. Lower class indices indicate stronger constraint. Points show 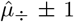 SE. **A:** CDS exons (1-10), [e]nhancers and [p]romoters; **B:** non-coding phastCons; **C:** `GPN-MSA` elements.

To quantify the influence of trinucleotide composition in exonic mutation rates, we recomputed 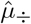 after filtering out CpG triplets. Ratios approach 1.0, as if the higher mutatability of exons is driven by the persistence of CpG sites as a result of first-order selection (Sved and Bird, 1990; Ségurel et al., 2014; Oman et al., 2022).

### Non-coding elements have lower predicted mutation rates than the neutral background

In stark contrast to protein-coding exons, both `GPN-MSA` and non-coding phastCons display lower mutation rates than the genomic background (Figure 1B-C). The depression of 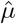 can reach 30% in non-coding phastCons and 40% in `GPN-MSA`, depending on mutation map. Masking CpG sites attenuates the effects–indicating that these elements are depleted in CpG content–but ratios remain *<* 1 across maps and constraint classes. To further explore the role of trinucleotide composition, we stratified 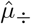 across the 64 motifs. Although the sharpest reductions occur at CpG sites (Figures A1–A2), ratios remain *<* 1 across most motifs. Only for a few does the Carlson map alone predict 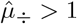, suggesting that the lower mutation rates attributed to these elements is not fully explained by their reduced CpG content compared to the rest of the genome. Notably, the relationship between 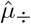 and predicted constraint is nearly monotonic for non-coding phastCons but clearly non-monotonic for `GPN-MSA` (Figure 1). Because phastCons are defined by phylogenetic conservation (Siepel et al., 2005), a negative correlation with mutation rate is expected whether the signal reflects biology or estimation bias. Conversely, `GPN-MSA`’s machine learning algorithm combines conservation scores from a vertebrate phylogeny with the 128-bp context surrounding each focal site. It is therefore difficult to pinpoint the reason behind the nonlinear relationship between 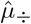 and predicted level of constraint in these elements. Another difference from non-coding phastCons is that `GPN-MSA` elements may overlap protein-coding exons (Methods). To investigate if this could explain their non-monotonic behavior, we computed 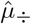 separately for `GPN-MSA` sites which overlap exons and for those which do not (Figure 2A). Ratios at non-exonic `GPN-MSA` sites retain their characteristic mean and shape, whereas in exonic sites they are close to 1.0 in the first four classes and peak near 1.5 before declining again. Therefore, the presence of exonic sites cannot explain the non-monotonic behavior of 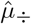 in `GPN-MSA` elements.

**Figure 2:**
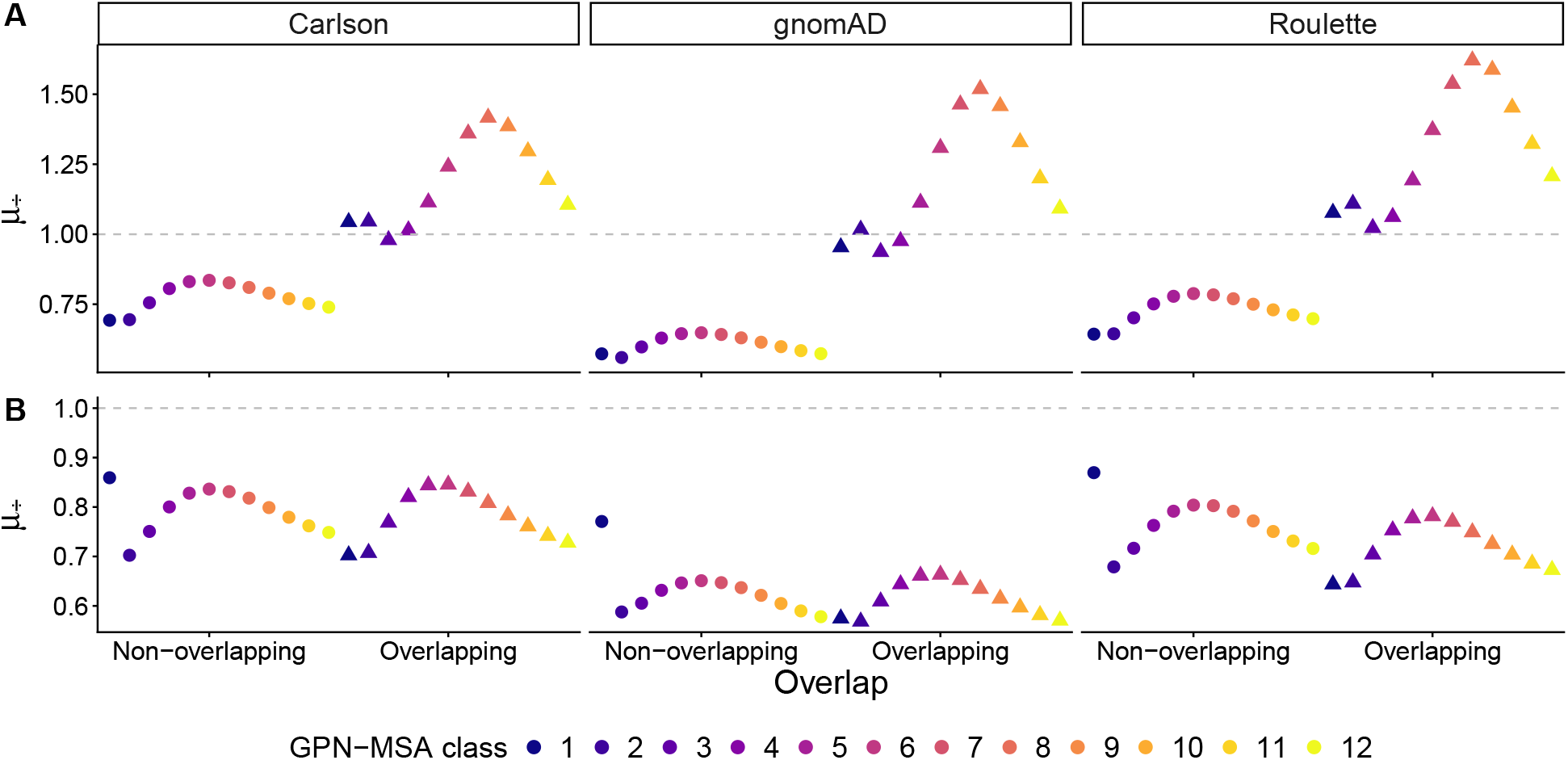
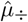 in `GPN-MSA` elements that do not overlap (circles) or overlap (triangles) other constrained sites. **A:** `GPN-MSA` overlapping protein-coding exons. Most sites which satisfy this condition belong to `GPN-MSA` classes 1-3 (strongest constraint). **B:** `GPN-MSA` overlapping non-coding phastCons have a 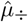 profile similar to those which do not. The higher 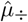 of `GPN-MSA` class 1 (darkest blue circle) is due to the presence of exonic sites with elevated mutation rates.

### The most likely direction of the estimation bias

What causes predicted mutation rates to be much lower in putatively constrained non-coding elements? A plausible concern is that the estimation of mutation rates using polymorhphism data is biased by background selection (BGS, Charlesworth et al., 1993). This possibility arises because both Carlson and Roulette use the spatial distribution of ERVs as a proxy for mutation rate variation on the scale of dozens of kb, on top of which they fit models of single-nucleotide resolution. The rationale behind approximating large-scale mutation rates by local ERV density is that these variants are young (mean age *≈* 50 generations, Fu et al., 2013) and therefore have been less affected by drift and selection, providing a relatively direct record of mutational processes along the genome (Carlson et al., 2018; Seplyarskiy et al., 2023). However, the focus on ERVs may not fully eliminate the influence of BGS, as it distorts the frequency spectrum (Cvijović et al., 2018) and may artificially depress 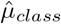 and 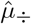.

That both maps nevertheless predict higher mutation rates in exons (where BGS is sharp) compared to the genome-wide background suggests that mutation-rate estimation using ERVs is not strongly confounded by BGS. Still, to further investigate the potential impact of BGS on our results, we focused on mutation rate differences at very short scales. We divided the genome in non-overlapping 1 kb windows, retaining only those harboring a single constraint class. This filter amplifies BGS differences among classes (quantified by their average *B*-value, see Methods), which are otherwise diluted by regions where sites from multiple classes co-occur. We then computed window-specific 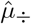, with 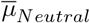 estimated from putatively neutral sites in the same windows as these isolated elements. We found no apparent trend between 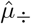 and *B*-values, and the overall patterns remain broadly consistent with genome-wide results (Figure 3). Moreover, because ERV-derived baseline mutation rates vary at *≈* 50 kb, they are constant within each of the 1 kb windows. Therefore, ratios *<* 1 at such fine scale cannot be explained by biases in the estimation of *µ* from ERV density, which would impact constrained and neutral sites equally. We conclude that 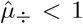 across non-coding elements is unlikely to result from BGS-induced errors in the mutation maps.

**Figure 3:**
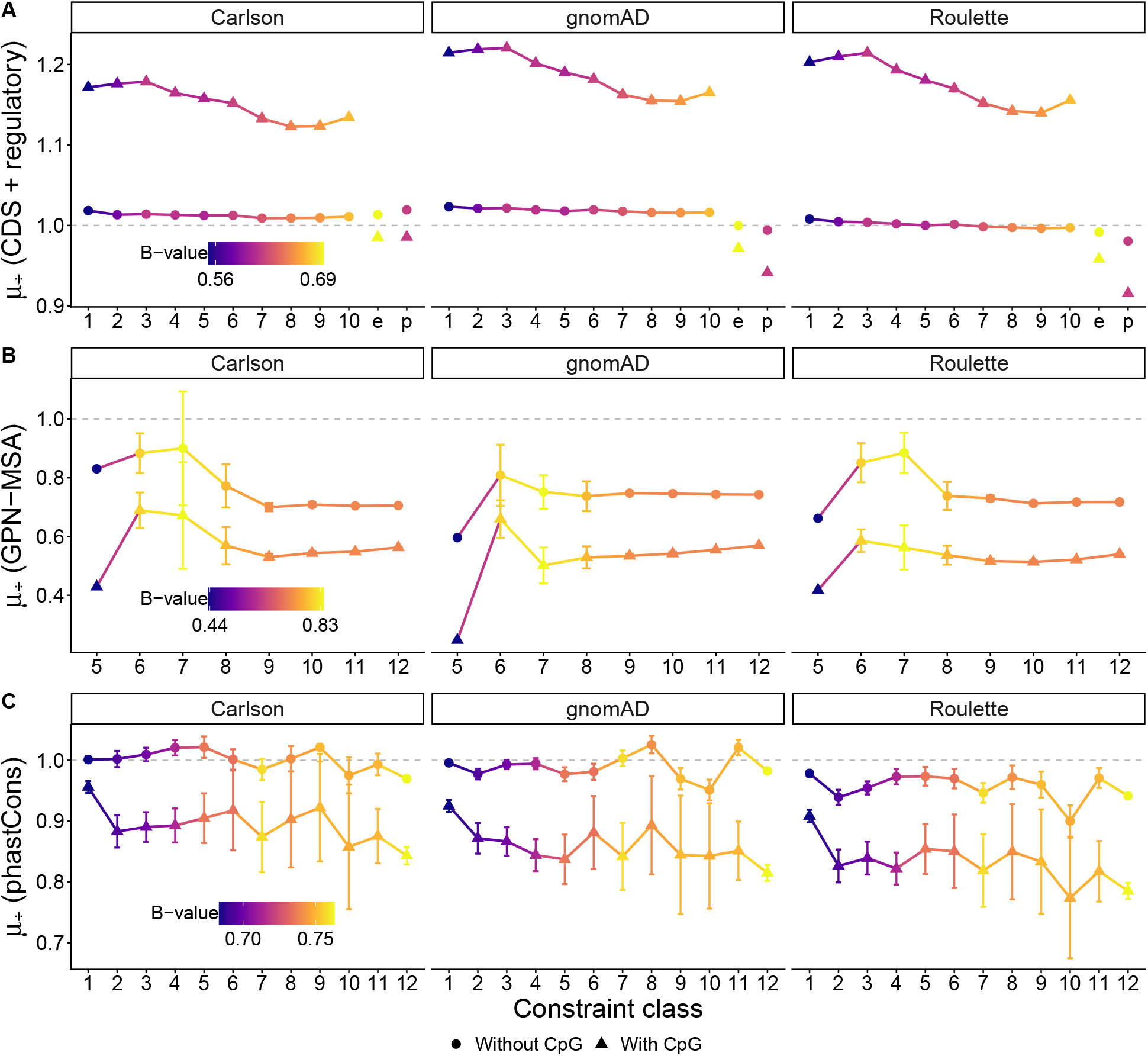
Predicted mutation rates differ in putatively constrained and nearby neutral sites. Points show median 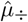 across 1 kb windows holding only one class of element (*±* 1 SE). **A:** CDS exons (1-10), [e]nhancers and [p]romoters. **B:** `GPN-MSA` elements, where 1 kb windows exclusively holding sites from classes 1-4 are absent. **C:** non-coding phastCons, where filtering out CpG sites brings ratios close to 1.0, indicating that the decreased mutability attributed to these isolated elements (which are very short, Figure A4) is mostly due to lower CpG content. A non-exclusive explanation is that the accuracy of phylogenetic scores varies with the fragmentation of synteny blocks in the multi-species alignment.

An alternative explanation is that the identification of constrained sites from sequence conservation has been biased by mutation rate variation. Specifically, sites with intrinsically low mutation rates may appear artificially constrained because they accumulate fewer substitutions across the phylogenetic tree. Consistent with this possibility, if non-coding elements were functionally constrained, selection would maintain hyper-mutable motifs as observed in CDS exons. Instead, non-coding elements appear depleted of CpG sites (Figures 1, 3), suggesting their identification has been biased by the simplifying assumption of constant mutation rates along the genome. Although there is convincing evidence of selection in non-coding phastCons as a whole (Di et al., 2025), the inferred strength of selection on classes beyond the first is weak, with mean 2*N*_*e*_*s ≈* 5 or less, so that some phastCons annotations may represent false positives. Similar arguments apply to `GPN-MSA`, which draws most of its information from local substitution rates on a vertebrate phylogeny (Benegas et al., 2025a). Hence, the negative correlation between mutation rates and non-coding constraint scores likely reflects over-identification of selection targets from sequence conservation.

### Accounting for mutation rate variation mitigates biases in constraint scores

During the preparation of these manuscript, Ye et al. (2026) developed `GPN-Star`, an improved version of `GPN-MSA` that introduces two important novelties. First, `GPN-Star` provides scores at three phylogenetic depths, with substitution rates estimated from alignments of vertebrates, mammals, or primates. As the authors show, this choice impacts the accuracy of `GPN-Star` in predicting the effect of different types of variants in humans: the vertebrate phylogeny is more informative about amino-acid changing mutations whereas the primate phylogeny is more informative about regulatory mutations, likely due to functional turnover. Second, to mitigate the confounding of mutation rate variation, `GPN-Star` is calibrated using the Roulette map. This procedure substantially decreases the correlation between `GPN-Star` scores and Roulette rates across chromosome 22 (held-out during model training), underscoring the importance of accounting for mutation rate variation in quantifying constraint (see Supplement in Ye et al., 2026).

However, diminishing correlations with Roulette rates across the entire spectrum of constraint does not imply that `GPN-Star` becomes fully independent of mutation rate variation. One possibility is that the bias persists in sites predicted to be under strongest constraint but is diluted by chromosome-wide comparisons. In this regard, Loay et al. (2026) argue that variant effect predictor models which correctly account for mutation rate variation suffer from a power imbalance whereby they can more easily detect depletion of substitutions in sites with higher mutation rates. This allows these sites to receive more extreme conservation scores, generating a spurious positive correlation with mutation rates. To investigate whether `GPN-Star` exhibits residual biases, we divided the top 30% scores across the genome into 12 equally-sized classes (as we did for `GPN-MSA`) and estimated *µ*_*÷*_ separately for each class and phylogenetic depth.

Overall, the genome-wide relationship between 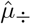 and `GPN-Star` scores (Figure 4A) is substantially improved relative to `GPN-MSA` (Figure 1C), especially for non-CpG sites (Figure A3). Nevertheless, 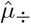 remains appreciably *<* 1 in the most constraint classes, with the magnitude of the bias depending on phylogenetic depth. This suggests that, despite calibration with the Roulette map, the deep-learning model may still capture residual variation in mutation rates. Notably, the downward bias in the most constrained classes goes in the opposite direction than predicted by Loay et al. (2026) for methods that correct for local mutation rates. One possible explanation is overfitting from scoring sites that were also used during model training.

**Figure 4:**
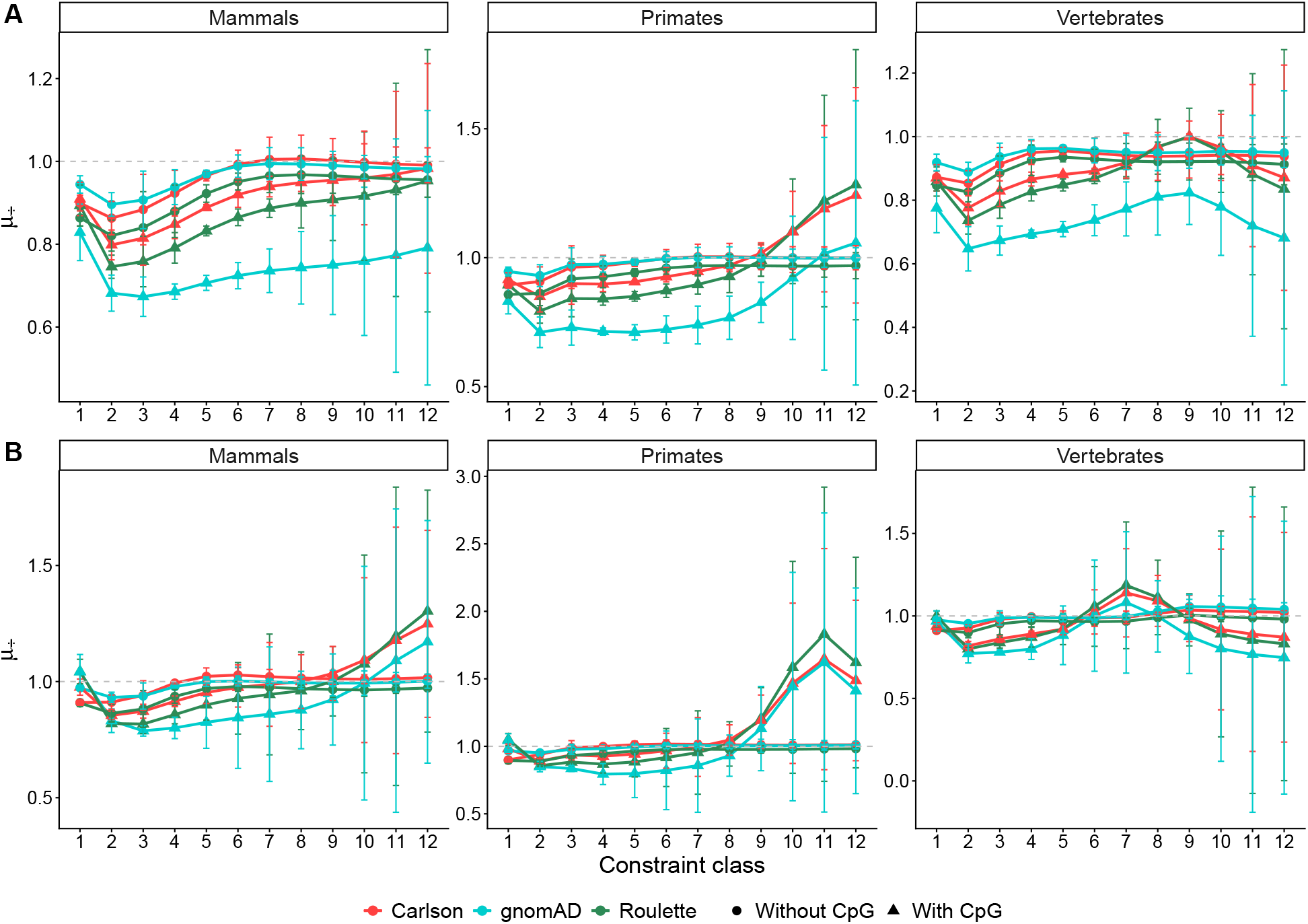
Despite calibration with the Roulette map, predicted mutation rates differ within and outside `GPN-Star` elements (top 30% scores split into 12 equal-size classes). Points show 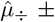 1 SE across genome-wide data (**A**) or chromosome 22 (**B**), which was held-out during model training. The SE correlates negatively with predicted constraint, indicating that the variance in mutation rates increases in collections of sites experiencing weaker selection.

The genome-wide trends are broadly recapitulated on chromosome 22, which was held out during training, although the bias is much weaker (Figure 4B). In the primate phylogeny, non-CpG sites show only a mild depression in mutation rates in the first few classes, after which 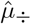 quickly approaches 1.0. These results are encouraging. They suggest that accounting for mutation rate variation can yield more reliable estimates of selective constraint, although there is still room for improvement in the most constrained sites. An alternative interpretation is selective optimization of local mutation rates. However, since estimation biases remain plausible, and since mechanisms that could reduce mutation rates in non-coding elements remain speculative, future work should prioritize mitigating potential biases before evoking adaptive mutation landscapes.

## Discussion

If *de novo* mutation rates genuinely differed between neutral and constrained sites, some foundational results in evolutionary biology would need revisiting. Here, we found that predicted mutation rates are depressed at putatively constrained non-coding sites in humans. Although this pattern is consistent with adaptive mutation landscapes, instead it likely reflects biases in the identification of selective constraint from mutation rate variation. Specifically, we postulate that phylogenetic scores have mis-classified sites with intrinsically low mutation rates as selectively constrained. A corollary of this result is that mutation landscapes are at least partially conserved across deeply diverged taxa, even at fine spatial scales (see also Beichman et al., 2023). Our findings complement a recent study by Loay et al. (2026), which showed that variant effect prediction within protein-coding genes is likewise biased by mutation rate variation. We further demonstrate that assessing the accuracy of constraint scores requires particular attention to outlier sites because genome-wide correlations can dilute biases concentrated in the regions of actual interest.

Selection on regulatory elements has driven key phenotypic changes during human evolution (Fraser, 2013; Hansen et al., 2024), motivating efforts to annotate the non-coding genome. Recent advances have opened new avenues for estimating site-specific levels of constraint from phylogenetic conservation (e.g. Benegas et al., 2025b,a; Avsec et al., 2026), but these approaches remain vulnerable to the confounding of mutation rate variation. Recognizing this limitation, `GPN-Star` explicitly incorporates local mutation rates into their predictions (Ye et al., 2026), leading to substantial improvements, yet biases appear to persist among sites experiencing the strongest constraint (Figure 4). In summary, we echo the concerns raised by McHale et al. (2026), who evaluated the performance of constraint metrics which are based on within-human diversity (Dukler et al., 2022; Halldorsson et al., 2022; Chen et al., 2024) rather than long-term conservation. The emerging picture from these benchmarking studies is that without a comprehensive account of the genomic features shaping patterns of variation, the identification of selection targets along the genome is compromised. We therefore suggest caution when interpreting results from the available non-coding annotations. Considerable uncertainty remains about their accuracy, and new methods should be designed carefully to avoid absorbing spurious signals.

Regardless of its underlying cause, our finding that predicted mutation rates are lower at putatively constrained non-coding sites has broad implications for human evolutionary genetics. Many studies rely on accurate mutation maps and on the identification of genomic regions subject to selection; systematic biases in either can consequently propagate into downstream inferences of evolutionary processes.

## Methods

### Preparing the data

Throughout, we used genomic maps and annotations in genome build GRCh38. We considered sites not included in each annotation set as belonging to the putatively neutral background. The processing of CDS, regulatory, and non-coding phastCons elements follows Barroso et al. (2026).

#### CDS elements

We downloaded exon data from the consensus coding sequence (CCDS) category in Ensembl release 114 (https://ftp.ensembl.org/pub/release-114/). From these, we removed sites which were not translated in any transcript, leaving only protein-coding sequence (comprising 33 Mb, 189,000 elements). We assigned coding (CDS) exons to genes using Ensembl gene identifiers and partitioned them into deciles of constraint according to gene *sh* scores from Zeng et al. (2024). (Genes not contemplated by Zeng et al. (2024) were excluded from the analyses). The estimated strength of selection decreases from class 1 to class 10 (see also Barroso et al., 2026).

#### Regulatory elements

We obtained annotations for promoters (29 Mb, 78,000 elements) and enhancers (239 Mb, 610,000 elements) after Di et al. (2025), who assembled these elements by combining promoter and enhancer-associated chromatin states inferred by Vu and Ernst (2022) using chromHMM (Ernst and Kellis, 2012). Promoter elements comprised chromatin states PromF1–PromF7 (flanking promoters) and enhancers comprised states EnhA1–EnhA20 (active enhancers) from Vu and Ernst (2022).

#### Non-coding phastCons elements

We obtained phastCons annotations after Di et al. (2025), who divided genome-wide phastCons conservation scores (17-way scores among primates, Siepel et al., 2005) into bins representing ~ 5% quantiles. They then intersected each bin with the set of candidate functional non-coding sites, defined as non-coding sites that were not classified as heterochromatin or quiescent chromatin states (regions with weak epigenetic marking) by Vu and Ernst (2022), and removed sites which did not pass the 1000 Genomes Project strict mask. These elements include the chromatin classes grouped into promoter and enhancer categories, as well as other regulatory elements and transcribed non-coding regions (introns, untranslated regions). In our analyses, we used the top 12 non-coding phastCons bins (756 Mb, 237 million elements), which contain the *≈* 60% highest phastCons scores. The predicted strength of selection decreases from class 1 to class 12.

#### GPN-MSA elements

We downloaded `GPN-MSA` scores from https://huggingface.co/datasets/songlab/gpn-msa-hg38-scores. These files contained constraint scores from the HG38 reference nucleotide to the three possible alternate nucleotides (Benegas et al., 2025a), which we summarized by the per-site average score. To mirror the analyses of phastCons elements, we defined the top 30% scores along the genome as putatively constrained sites. We divided them into 12 equally-sized classes, but unlike in the non-coding phastCons annotations, here we kept sites overlapping CDS exons. The predicted strength of selection decreases from class 1 to class 12.

#### GNP-Star elements

We downloaded `GPN-Star` scores from https://huggingface.co/datasets/songlab/gpn-star-scores. These files contained constraint scores (available for primates, mammals or vertebrates phylogenies) from the HG38 reference nucleotide to the three possible alternate nucleotides (Ye et al., 2026), which we summarized by the per-site average score. To mirror the analyses of phastCons elements, we defined the top 30% scores along the genome as putatively constrained sites. We then further divided these sites into 12 equally-sized classes, but unlike in the non-coding phastCons annotations, here we kept sites overlapping CDS exons. The predicted strength of selection decreases from class 1 to class 12.

#### Mutation maps

We obtained site-resolution Carlson (Carlson et al., 2018), gnomAD (Karczewski et al., 2020) and Roulette (Seplyarskiy et al., 2023) mutation rates from https://genetics.bwh.harvard.edu/downloads/Vova/Roulette/. These `VCF` files contained mutation rates from the HG38 reference nucleotide to the three possible alternate nucleotides. To calculate a site-resolution average rate, we took the sum of rates at each site and applied the scaling factors provided at https://github.com/vseplyarskiy/Roulette.

### Estimating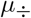

We intersected annotation groups and mutation maps to estimate mutation rates within and outside putatively constrained sites with single-nucleotide resolution. We first estimated *µ*_*÷*_ genome-wide (Figure 1) as the ratio of mutation rates within each constraint class and the neutral genomic background (i.e., sites not belonging to any constraint class of the focal annotation group), 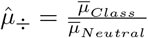.

We then divided the genome into non-overlapping 1 kb windows, and discarded any window that contained constrained sites from more than a single class. We performed this filtering separately for each annotation group. We then merged these data with 1-kb resolution *B*-value maps predicted with `moments++` (Barroso and Ragsdale, 2026) which represent the relative reduction in diversity caused by background selection in humans (Barroso et al., 2026). When estimating window-specific *µ*_*÷*_ (Figure 3), we considered only putatively constrained and neutral sites belonging to each window.

We also stratified *µ*_*÷*_ by trinucleotide context using the two sites adjacent to the focal site (defined by the reference alleles in the Roulette `VCF` files, Seplyarskiy et al., 2023). We used these stratified data to filter out CpG sites (Figures 1, 3) as well as to average mutation rates 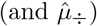 per triplet (Figures A1–A3).

## Data availability

Scripts we used are available in https://github.com/gvbarroso/mut_rates_func. The repository contains the summarized representations of the data (constraint scores and mutation rates) required to reproduce figures and statistical analyses.

## Acknowledgments

We thank David Castellano, Fanny Pouyet and members of the Ragsdale lab for helpful discussions. Computational resources were generously provided through UW-Madison’s Center for High Throughput Computing. This work was supported by NIH Award R35GM154962 to APR.

## Extended Data Figures

**Figure A1:**
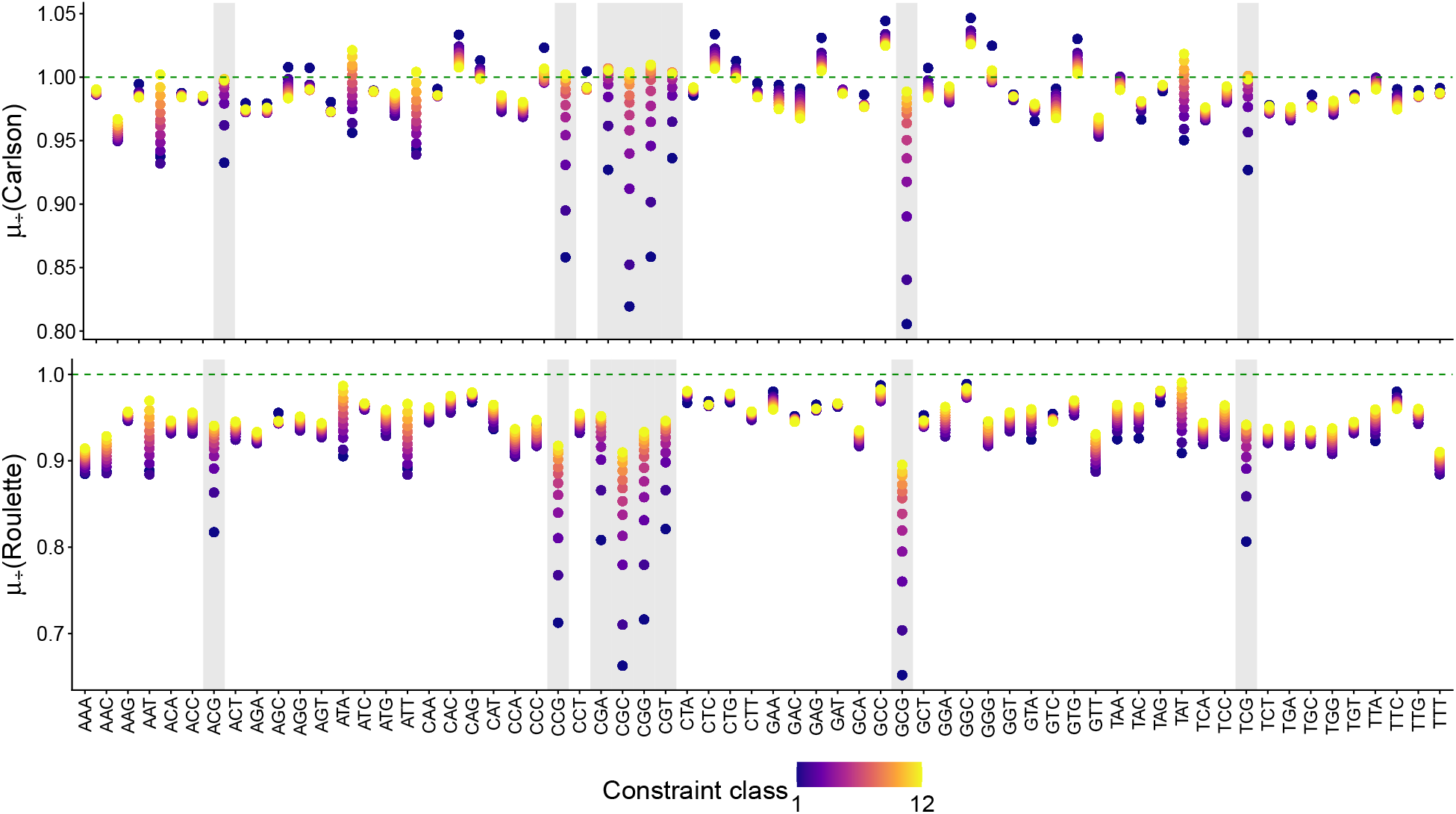
Ratios computed for phastCons elements and stratified by triplet context. By construction,the gnomAD mutation map stands on top of the 1 line (dashed green).

**Figure A2:**
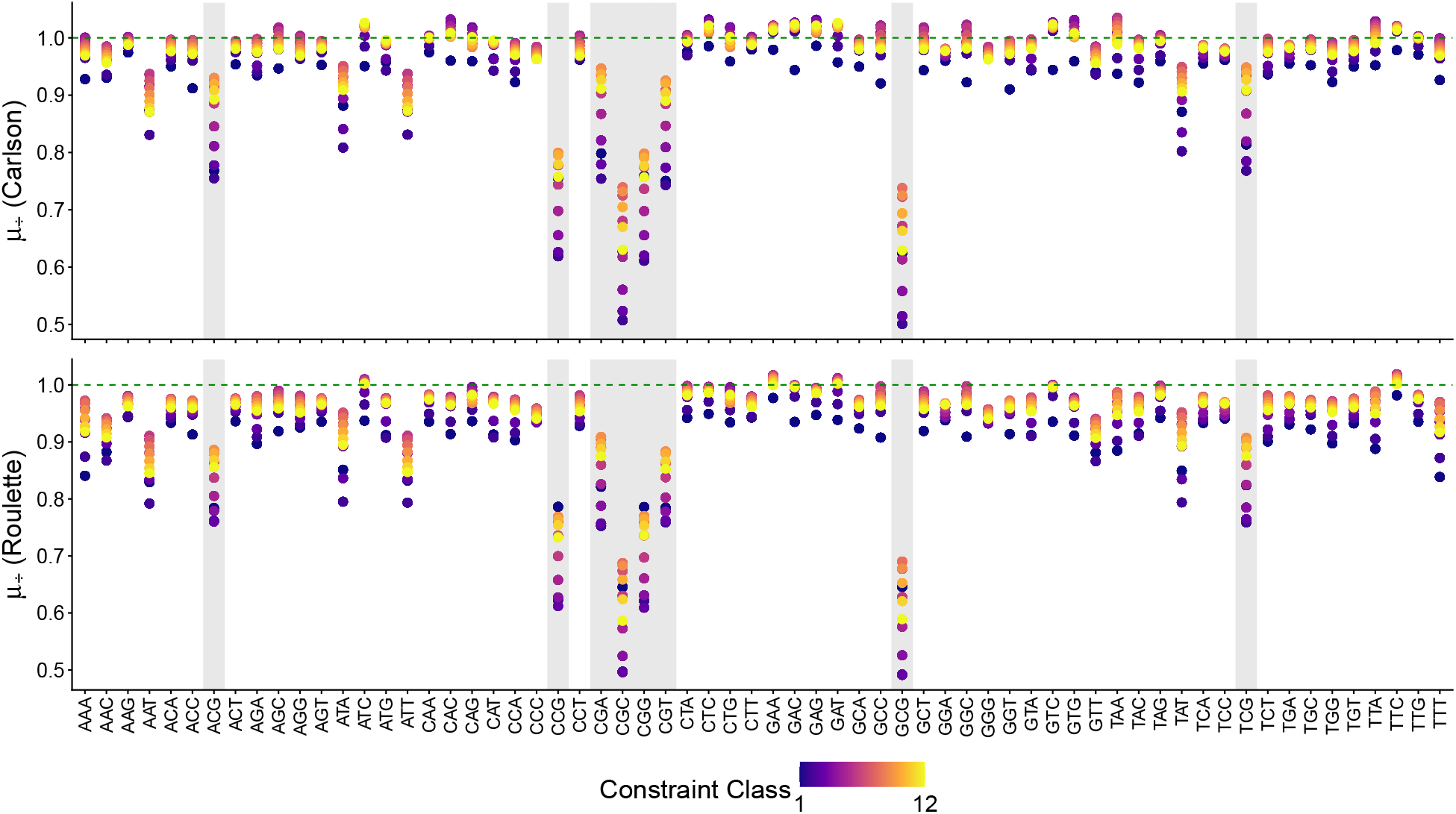
Ratios computed for `GPN-MSA` elements and stratified by triplet context. By construction, the gnomAD mutation map stands on top of the 1 line (dashed green).

**Figure A3:**
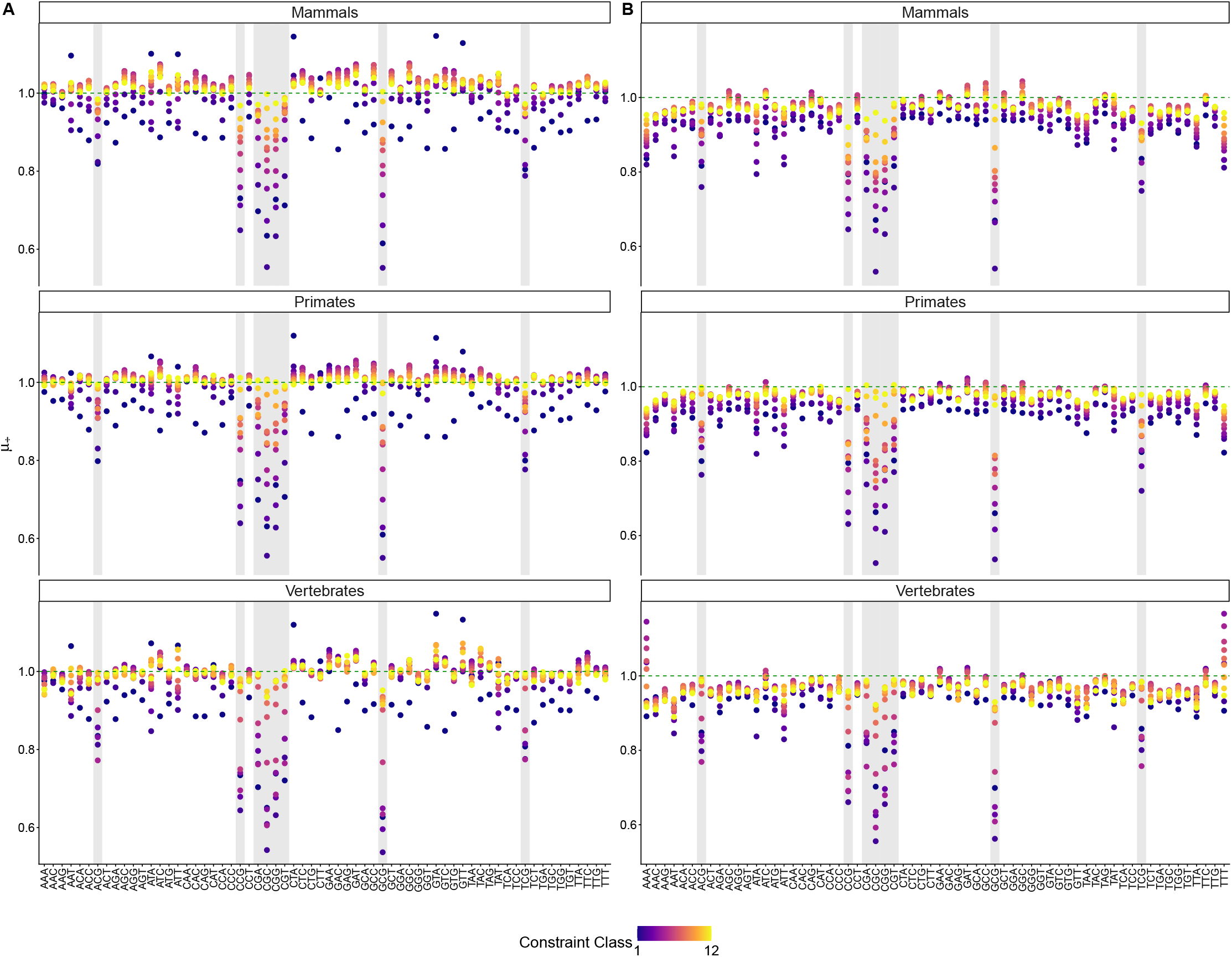
Genome-wide ratios computed for `GPN-Star` elements and stratified by triplet context. **A:** Carlson map. **B:** Roulette map. Each panel corresponds to a different phylogenetic depth. By construction, the gnomAD mutation map stands on top of the 1 line (dashed green).

**Figure A4:**
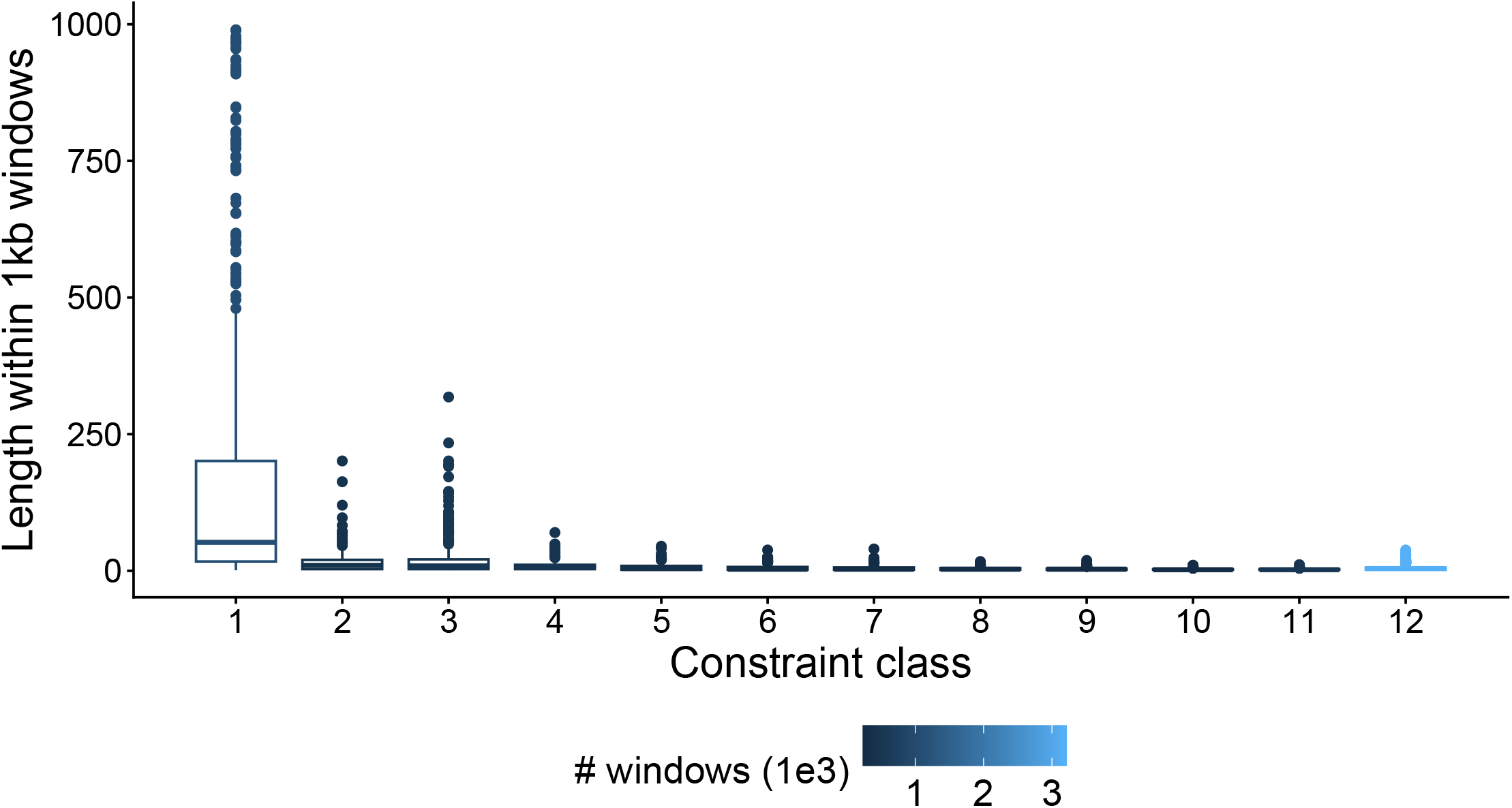
Distributions of total lengths of each non-coding phastCons class within 1 kb windows. Except for the most constrained class, these elements are very short. Color key shows number of 1 kb windows (in units of thousands) harboring only sites of a given class.

